# RNAbridge: a database of extended non-canonical helices

**DOI:** 10.64898/2026.09.15.751775

**Authors:** Damian Zakrzewski, Maciej Antczak, Tomasz Zok

## Abstract

RNA function is dictated by 3D architecture. Although 2D structural models based on canonical Watson-Crick base pairs are widely used, they often fail to capture the non-canonical interactions, tertiary contacts, and coaxial stacking important for biological activity. We have developed RNAbridge, a comprehensive database and web application that systematically identifies, quantifies, and visualizes extended non-canonical helices and multi-way junctions (“non-canonical” denoting the interruptions between canonical stems rather than non-canonical pairing within them). Using a geometry- and stacking-based pipeline, we analyzed the Protein Data Bank and compiled a catalog of 153,148 structural elements, with a representative view of 7,453 elements. In this view, 4,998 multi-segment extended helices and junction arrangements have measurable bend angles, and 2,455 are single-segment canonical stem-loops. RNAbridge includes a user-friendly interface with interactive filters, synchronized 2D and 3D visualizations, and detailed structural data, including helical bend angles and stacking paths. By connecting simplified 2D topologies with complex 3D structures, RNAbridge serves as a valuable resource for structural biologists and lays the foundation for future machine learning applications in RNA structure prediction. The database, available at https://rnabridge.cs.put.poznan.pl/, is automatically updated once a week. All counts reported here refer to the November 8, 2026 release.

## 1 Introduction

RNA molecules adopt sophisticated folds that enable them to function as biological catalysts (ribozymes), molecular sensors (riboswitches), and key elements of the protein-translation machinery. Computational studies of large RNAs typically operate on simplified 2D secondary-structure models built from canonical Watson-Crick and G-U wobble base pairs [1]. This restriction is what makes folding prediction and scaffold mapping tractable for long sequences, but it comes at the cost of biological accuracy. The functional properties of RNA depend on non-canonical base pairs, such as Hoogsteen interactions, and tertiary contacts, including base triples, pseudoknots, and A-minor motifs [2–4]. Simplified 2D models often fail to capture the structural complexity and conformational dynamics underlying catalytic activity and regulatory specificity because they overlook these non-standard interactions, along with the stabilizing effect of metal ions [5]. Tools for scoring secondary-structure models against non-canonical base pairs exist [6], but they do not treat stacking-connected helices as objects.

The structural complexity of RNA largely arises from loops, including hairpins, internal loops, and junctions, rather than from stems. These regions frequently enable coaxial stacking [7], resulting in elongated, non-canonical helices formed from canonical stems connected by tertiary interactions. The interaction between structured stems and dynamic loops converts a linear base sequence into a functional, three-dimensional molecular machine [8].

A canonical stem consists of consecutive Watson-Crick C-G and A-U pairs and wobble G-U pairs. An extended non-canonical helix consists of two or more canonical stems connected by continuous stacking paths across bulges, internal loops, or n-way junctions, provided they satisfy the geometric criteria described below. In this context, “non-canonical” refers to the interruptions between stems rather than non-canonical pairing within the stems. We classify extended helices by the number of constituent stems.

RNAbridge is a database of extended non-canonical helices and coaxially stacked multi-way junctions. Each entry comprises canonical stems connected by loops with defined interactions, enabling the structure to be analyzed as a single unit. The catalog also includes selected single-segment stem-loops whose terminal hairpins satisfy the stacking-continuity criterion (Section 3.4). The database includes elongated non-canonical helices formed by coaxially stacked stems inter-spersed with internal loops or bulges. It also catalogs arrangements from multiloop junctions and identifies coaxially stacked stems within these regions. Whereas the RNA 3D Motif Atlas [9] groups recurring local geometries into discrete classes, RNAbridge extracts stacking-connected helical and junctional elements and describes their topology and geometry. These approaches are complementary. RNAbridge offers a user-friendly interface, regular automatic updates, interactive visualizations, and downloadable results.

The available related resources divide up the space of structural arrangements either locally (SCOR, RNA 3D Motif Atlas, RNA CoSSMos, RNA Bricks, RNA FRABASE, RNAMotifScanX) [9–15], on the basis of topology (RAG, CRW) [16, 17], according to interaction type (NCIR, InterRNA, PseudoBase) [18–20], or for use in machine learning (RNANet, RNAsolo, RNA3DB) [21–24]. Large-scale prediction benchmarks such as CASP [25] and structural challenges such as RNA-Puzzles [26] and its associated computational toolkit [27] highlight the ongoing difficulties in modeling complex RNA folds [28]. Junction geometry can be explored using resources such as RNAJunction and RNAloops, which support searches by inter-helical angle [29, 30]. However, none of these treats a coaxially stacked, internally interrupted helix as an object with its own identifier, statistics, or rendering.

From a user perspective, all 3D motif databases store sequence and tertiary structure, can be searched by PDB ID and sequence, and offer tertiary structure downloads. Most handle all motif types, while RNAJunction and RNAloops focus only on multi-branch loops. Secondary structures are stored and can be queried by all 3D motif databases considered except the RNA 3D Motif Atlas and RNAJunction. Angular data is offered only by RNA FRABASE and RNAloops. Motif topology search is offered by FRABASE, RNA Bricks, and RNAloops, and RAG offers only graph-based features. All databases except RNAJunction are regularly updated. All currently existing databases focus on independent 3D motifs. However, RNAbridge treats helices that run through junctions and non-canonical regions as specific objects equipped with coaxial stacking information.

RNAbridge allows users to search for experimentally observed coaxial arrangements around n-way junctions and inspect their sequence and structural context. These arrangements may provide templates or constraints for RNA 3D structure modeling.

## 2 Results and Discussion

### 2.1 Web application

The RNAbridge web interface provides an interactive environment to explore and analyze RNA extended non-canonical helices and multi-way junctions. Users can search and filter datasets using criteria such as PDB identifier, experimental method, nucleotide sequence patterns, type, number of segments, global bend angle range, and nucleotide count, with the option to restrict results to a representative view labeled “non-redundant” in the interface (Figure 1A). An interactive statistics dashboard, a histogram of bend angle distributions, and a pie chart of element composition summarize the search results. Both the result table and these summaries are recalculated for the active selection, so they reflect the current filters rather than the full catalog. All chart elements act as active filters, enabling users to refine results directly through the visualization.

**Figure 1:**
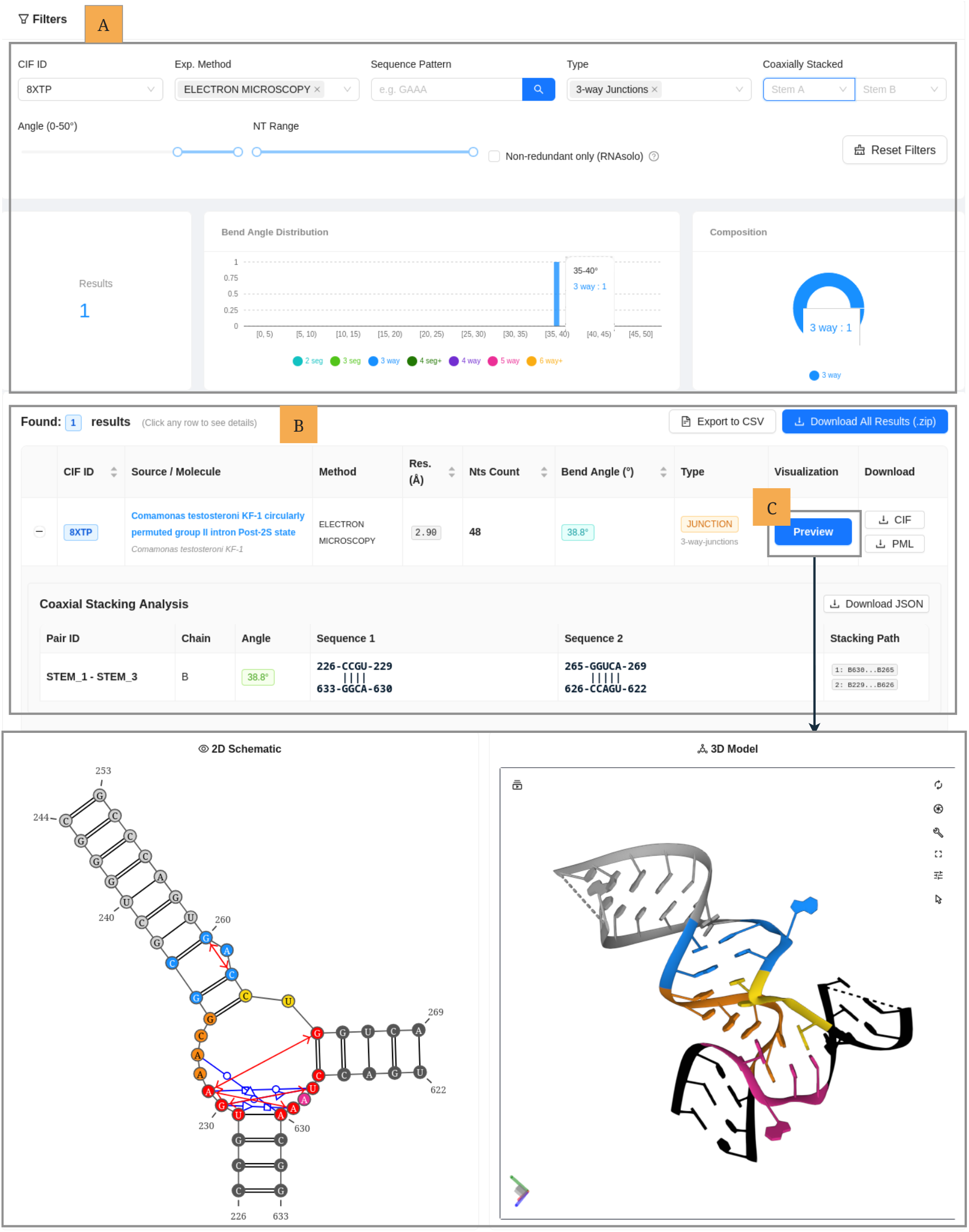
RNAbridge overview. (A) Available filters, (B) a row with search results, (C) 2D and 3D visualizations.

A structured table displays the filtered records alongside PDB metadata, including molecule name, source organism, experimental method, and resolution, as well as structural metrics (Figure 1B). Users can expand each record to inspect its detailed structural breakdown. For helices, the view displays individual segments of stems, bulges, and hairpins, together with their stacking paths, local bend angles at the connectors between stems, and a comparison of two-dimensional and three-dimensional nucleotide counts. For junctions, the view displays identified coaxial stem pairs, relative inter-stem angles, and schematic diagrams of stacking bridges. A synchronized side-by-side layout visualizes structure by combining a two-dimensional topology schematic with an interactive three-dimensional model powered by Mol* [31] (Figure 1C). Consistent colors across both views help identify individual structural parts. Users can also export filtered results to CSV format and batch-download mmCIF coordinate files and PyMOL [32] scripts bundled in a single ZIP archive.

### 2.2 Database content

The full catalog contains 153,148 structural elements extracted from 5,220 RNA-containing PDB entries. These elements are divided into three classes: 49,226 single-segment canonical stem-loops, 61,994 multi-segment extended helices, and 41,928 junction-based arrangements. Single-segment entries are canonical stem-loops with a terminal hairpin that satisfies the stacking-continuity criterion (Section 3.4). Therefore, they do not represent a complete census of all canonical stem-loops in the PDB. Three-way junctions are the most common junction class (20,543), followed by four-way junctions (14,777), five-way junctions (2,873), and junctions of order six or more (3,735). The latter group comprises 1,408 six-way, 2,325 seven-way, and 2 junctions of order eight or higher. No finer breakdown above seven is reported here.

Entries are classified topologically: helices are categorized by the number of constituent stems (up to seven in the current release), and junctions by their order. Descriptors including global and local bend angles, stacking paths, nucleotide counts, constituent-stem sequences, and non-canonical base-pair composition support the retrieval of structurally related substructures. The catalog inherits the composition of the Protein Data Bank and is therefore non-uniform.

A few frequently solved systems dominate the extracted elements: ribosome structures and their subunits contribute a disproportionate share, whereas most RNA families are represented by only a few structures. The frequency of an arrangement in the full catalog reflects the experimental solve rate of the corresponding RNA, not its biological prevalence or functional importance.

Because source structures are highly redundant, the database provides a representative view, labeled “non-redundant” in the interface. Entries are retained when their RNA chains belong to a representative chain set in the BGSU non-redundant list [33], accessed through the RNAsolo2 API [24] (BGSU 4.64, November 5, 2026). Selection therefore operates on RNA chains rather Angle / Type Multi-segment Helices Junction-based Arrangements Total than whole PDB entries, without additional experimental-method or resolution filtering. This view contains 7,453 elements from 1,212 PDB entries: 2,455 single-segment canonical stem-loops, 3,136 multi-segment extended helices, and 1,862 junction-based arrangements. This is a 20.5-fold reduction in element count relative to the full catalog. We use this view for the composition statistics below, while the full-catalog distributions and threshold sensitivity are reported in Tables S1, S3, and S4. Representative-source selection reduces repeated sampling but does not guarantee that local elements are non-redundant. Machine-learning applications additionally require sequence- or structure-aware train/test partitioning to limit data leakage.

We calculated bend angles for 4,998 multi-segment and junction elements in the representative view with *θ* ≤ 50° (Table 1). The 2,455 single-segment entries lack an inter-helical angle. The corresponding full-catalog count is 103,922 (Table S3). Within the accepted range, the distribution is unimodal and skewed toward low bend angles. It peaks in the [10°, 15°) bin (1,008 elements, 20.2%), and 62.0% of the 4,998 elements have bends below 20°. This distribution decreases toward the upper limit, with 59 elements (1.18%) in the [45°, 50°]interval.

**Table 1.** Bend-angle distribution of multi-segment helices (2-seg to 4-seg+) and junction-based arrangements (3-way to 6-way+) in the representative view, at the default cutoff (*θ* ≤ 50°). Helix values are global first-to-last stem angles; junction values are the smallest angles among their selected coaxial stem pairs. The view contains 7,453 elements, of which 4,998 have a measurable bend angle. The 2,455 single-segment canonical stem-loops are excluded because no inter-helical bend angle can be computed. Counts refer to the November 8, 2026 release.

| Angle / Type | Multi-segment Helices |  |  | Junction-based Arrangements |  |  |  | Total |
|---|---|---|---|---|---|---|---|---|
|  | 2-seg | 3-seg | 4-seg+ | 3-way | 4-way | 5-way | 6-way+ |  |
| [0°, 5°) | 215 | 7 | 4 | 44 | 83 | 22 | – | 375 |
| [5°, 10°) | 484 | 25 | 7 | 117 | 185 | 43 | – | 861 |
| [10°, 15°) | 598 | 33 | 2 | 194 | 149 | 17 | 15 | 1,008 |
| [15°, 20°) | 522 | 56 | 4 | 154 | 81 | 15 | 25 | 857 |
| [20°, 25°) | 394 | 43 | 4 | 108 | 69 | 11 | 18 | 647 |
| [25°, 30°) | 239 | 35 | 2 | 83 | 63 | 1 | 37 | 460 |
| [30°, 35°) | 189 | 44 | 6 | 50 | 47 | 7 | 14 | 357 |
| [35°, 40°) | 97 | 15 | 7 | 58 | 57 | 1 | 3 | 238 |
| [40°, 45°) | 49 | 23 | – | 34 | 26 | – | 4 | 136 |
| [45°, 50°] | 21 | 11 | – | 19 | 8 | – | – | 59 |
| Total | 2,808 | 292 | 36 | 861 | 768 | 117 | 116 | 4,998 |

Two-segment helices account for 2,808 of the 4,998 angle-analyzed elements (56.2%) and 89.5% of the 3,136 multi-segment helices. The six-way or higher junction class differs from the general trend: its modal bin is [25°, 30°) (37 of 116 elements, 31.9%).

Element size, measured as the number of unique nucleotides in each reported element (Tables S1 and S2), is concentrated at 10–29 nucleotides, accounting for 81.0% of elements in the representative view. Fewer than 1% contain 50 or more nucleotides. Of the 11 elements containing at least 60 nucleotides, 10 are multi-segment extended helices. The full catalog shows a similar concentration (Table S1).

### 2.3 Case studies

#### 2.3.1 Helix 44 of the small ribosomal subunit

Helix 44 is the longest and most conserved helix of the small ribosomal subunit RNA (16S rRNA in bacteria, 18S rRNA in eukaryotes) and has a direct role in the fidelity of translation initiation [34]. In the human structures (PDB IDs: 7QP6/7QP7 [35]; cryo-EM reconstructions at 4.7 Å and 3.7 Å, respectively), it corresponds approximately to nucleotides 1656–1812 of 18S rRNA. This region forms a long, nearly continuous A-form double helix that extends along the back of the 40S subunit body, from the solvent side to the decoding center at the interface.

RNAbridge identifies the regions 1722-1750 and 1784-1812 of the 7QP7 structure, chain A, as forming a five-segment extended helix (Figure S1). The first and third internal loops are simple pyrimidine-pyrimidine mismatches consisting of U1730-U1804 and C1740-C1794, respectively, and these interact in a cis Watson-Crick/Watson-Crick manner, similar to canonical base pairs. The second internal loop is symmetric and comprises a pair of inverted base pairs: first G1734 paired with A1800 in a trans Sugar/Hoogsteen configuration, then A1735 with G1799 in a trans Hoogsteen/Sugar configuration. Because of the way these base pairs are arranged, the stacking paths are altered so that G1734 stacks with G1799 and A1735 stacks with A1800. The last internal loop is a 3 *×* 3 loop closed by three pairs: G1743-A1791 in cis Sugar/Hoogsteen, G1744-G1789 in cis Sugar/Watson-Crick, and A1745-G1789 in cis Hoogsteen/Sugar arrangements. As a result, even though the loop is entirely composed of purines, the network of interactions causes the nucleobases to be oriented inward rather than projecting outward; this enables the entire five-segment element to appear as, and to be detected as, a single, extended helix that is almost like the A-form. Because a 3.7 Å map resolves the path of the duplex but not individual hydrogen bonds, the base-pair classes given here indicate probable geometry rather than exact hydrogen-bond patterns.

### 2.3.2 Three-way junction families

Twenty years ago, Lescoute and Westhof outlined the possible topologies of three-way junctions [36], noting that two helices generally adopt a coaxial configuration. At the same time, the third helix occupies different positions mainly depending on the lengths of the single-stranded linkers. They identified three possible configurations for the third helix and named them Families A, B, and C. These families remain a useful framework for describing three-way junctions. Junction-focused resources such as RNAJunction and RNAloops support geometric analysis and angle-based searches [29, 30]. RNAbridge complements them by letting users relate this family-level description to the underlying stacking and non-canonical interactions in synchronized 2D and 3D views, making a junction’s internal architecture directly inspectable.

As stated in the foundational study by Lescoute and Westhof, Family C is “the most fascinating one” because it occurs in various RNAs with different functions. This family is characterized by a long J31 strand which links the 5’ strand of P3 – the non-coaxially stacked helix – with the 3’ strand of P1 – the first helix. This extended J31 strand is thought to be highly structured. We examined the crystal structure of the specificity domain of RNase P RNA (PDB ID: 1NBS [37]), which is known to have a Family C junction [36]. Our analysis confirms this previously known information (Figure S2), but RNAbridge stands out for how easily it identifies the feature and how clearly it presents the results. In the 2D visualization, the J31 strand is shown in yellow, and it is clear that the five nucleotides within it form two base pairs: U175-A179 trans Watson-Crick/Watson-Crick and G176-A178 trans Sugar/Hoogsteen in a tight arrangement. The same yellow color in the 3D diagram shows how closely the nucleotides are packed together. This network of interactions enables P3 to bend toward P1 – the defining feature of Family C and one clearly illustrated by RNAbridge.

## 3 Materials and Methods

### 3.1 System architecture

The system runs on a virtual machine with Debian GNU/Linux 13 (Trixie), 8 GB of RAM, and a multi-core CPU. Docker containerizes the entire stack to guarantee consistent behavior across deployment environments, managing both the virtual machine provisioning and the Anaconda environment.

The backend uses Python with the FastAPI framework and stores data in PostgreSQL through the SQLAlchemy Object-Relational Mapping tool. The frontend is built with React, TypeScript, and the Ant Design component library. The computational pipeline for structural analysis is written in Python and uses RNApolis Annotator [38], the successor of the RNApdbee 2.0 annotation server [39], PyMOL [32] and X3DNA [40]. The system retrieves structural data from the RCSB Protein Data Bank [41]. For visualization, it renders two-dimensional secondary structures with VARNA (varna-tz) [42] and displays three-dimensional molecular graphics with Mol* [31].

The RNAbridge database implements an automated weekly synchronization mechanism with the RCSB PDB. During each weekly update cycle, the system fetches the current set of active RNA identifiers and scans local storage. Structures that have been deprecated or removed from the PDB are automatically purged—along with their associated local CIF files, JSON metadata, result directories, and database records. This process ensures that the local repository and database remain fully consistent with the primary repository.

The catalog includes structures regardless of the experimental method or resolution. The reported statistics pool all methods without method-specific weighting. We record the experimental method when available, which users can use as a filter. We label missing annotations as “Unknown.” When selecting structures for analysis, users should consider differences in coordinate precision and, for NMR, ensemble representation. Only the first model in each multi-model structure is analyzed, and alternate conformations are resolved by occupancy.

### 3.2 Processing pipeline

The workflow begins with data ingestion and preprocessing (Figure 2). In this stage, the pipeline normalizes mmCIF files, retrieves experimental metadata from the Protein Data Bank, and annotates the structures automatically. The annotator is run in pseudoknot-free decomposition mode: pseudoknot stems are reported separately and are not cataloged as independent stems. Pseudoknot characters retained in the loop structure are counted as loop nucleotides, and loops containing pseudoknot-associated nucleotides are not discarded. The helix and junction catalog therefore includes pseudoknot-associated junctions without treating pseudoknot stems as separate helical elements. It then builds high-speed lookup indices to catalog all canonical and non-canonical interactions, enabling efficient queries of the resulting structural network. Next, the pipeline classifies entries to identify and label fundamental structural elements, including hairpins, bulges, and N-way junctions.

**Figure 2:**
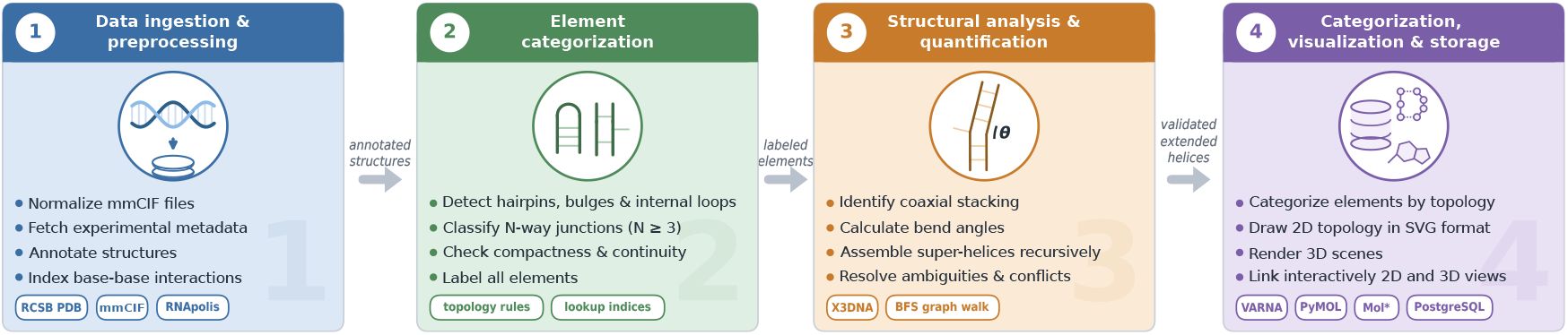
RNAbridge processing pipeline.

Structural analysis and quantification follow. The pipeline verifies coaxial stacking stability using breadth-first search pathfinding and measures 3D helical bend angles with X3DNA. It then recursively assembles detected elements into extended helices based on spatial orientation and geometric distance. Finally, the system categorizes the assembled architectures by topology – the same categories shown as types in the web interface – and renders them as interactive two-dimensional (SVG) and three-dimensional (PML/mmCIF) visualizations. It maps all biological metadata and hierarchical results into a searchable relational database for storage and downstream integration.

All results reported here were produced with the following configuration. Structures are annotated with RNApolis Annotator in pseudoknot-free decomposition mode (`annotator --decompose-pseudoknot-free`), with a ten-minute limit per structure. Structures that exceed it are skipped. The stacking graph is built directly from the annotator’s base-stacking interaction list and is treated as an unweighted, undirected graph. Helical axes are computed with X3DNA v2.4 using the `find_pair` and `analyze` routines. Two-dimensional diagrams are drawn with VARNA through varna-tz, three-dimensional scenes with PyMOL (open-source, version 3.1 or later), and interactive molecular views in the browser with Mol*. The bend-angle cutoff is a single parameter, set to 50° for all runs reported here. The counts obtained at the other thresholds are given in Table S4.

### 3.3 Bend angle

We characterized the three-dimensional geometry of the extracted RNA structural elements by calculating inter-helical bend angles from the central axes of their constituent stems. For each canonical RNA stem, we determined the local helical axis using the `find_pair` and `analyze` routines from X3DNA. To ensure consistent orientation, we oriented each axis vector from 5’ to 3’, based on the spatial arrangement of the C1’ atoms along the stem’s 5’ strand. We calculated the bend angle between two stems as *θ* = arccos(**u**_1_ ·**u**_2_), where **u**_1_ and **u**_2_ are the normalized helical axis vectors. By construction, *θ* ranges from 0° (parallel, co-oriented axes) to 180° (antiparallel axes). In practice, the assembly pipeline only accepts configurations with *θ* ≤ 50 °(see Helix extension and Junction analysis), so all reported angles fall within [0°, 50°]. We justify this cutoff and assess the sensitivity of the results to its value in Section 3.3.1.

We adapted the measurement strategy to each element’s structural context. For multi-segment helices, we quantified the overall deformation caused by structural interruptions, such as internal loops or bulges. We calculated a global bend angle from the axis vectors of the first and last flanking stems of the contiguous extended helix to represent the cumulative spatial deflection of all intervening elements.

For branched topologies, such as multi-arm junctions, we determined helical axes for all radiating stems and calculated bend angles for every pairwise combination. For a four-way junction, this approach yields six pairwise inter-helical angles, which describe the junction core and help identify coaxial stacking interactions. The summary angle reported for each junction is the smallest angle among its selected coaxial pairs. The global angle for a helix is measured between its first and last stems.

#### 3.3.1 Choice of the bend-angle threshold

Stacking continuity identifies candidate connections independently of the angular criterion (Section 3.4). The 50° cutoff is a separate geometric inclusion rule. We chose this permissive boundary to retain moderately bent arrangements. However, it is not an experimentally established boundary between coaxial and non-coaxial structures.

To assess sensitivity to this choice, we repeated the assembly at 50°, 70°, and 90° (Table S4). Because the threshold affects assembly rather than post hoc filtering, we re-derived the retained set for each value. The full-catalog count of angle-analysed elements increased from 103,922 at 50° to 105,922 at 70° (+1.9%), and to 106,167 at 90° (+0.23% relative to 70° and +2.2% relative to 50°). Most of this increase occurred in junction classes: 3-way and 4-way counts increased by 6.2% and 5.2%, respectively, between 50° and 70°, while the 2-segment count decreased from 56,236 to 55,708. While the total counts changed little as the cutoff relaxed, the individual assemblies and category assignments varied.

### 3.4 Helix extension

The assembly of n-segment helices relies on a two-stage validation procedure applied to stem-connecting connectors, such as internal loops and bulges. In the first stage, the algorithm assesses each connector by its stacking status, determined through a breadth-first search traversal of the nucleotide stacking graph. Stacking interactions are identified by RNApolis Annotator, using base-center distance and angular criteria. An element qualifies as a valid connector only when full stacking continuity is preserved across both flanking stems. Internal loops require two node-disjoint stacking paths connecting terminal nucleotides on opposite sides of the duplex, whereas bulges require a path across the interrupting strand.

Terminal hairpins are validated separately because they close a helix rather than connect two stems. A hairpin is accepted if the loop-restricted stacking graph contains a continuous path between the nucleotides of its closing base pair. This path may include unpaired or non-canonically paired loop nucleotides. If the test fails, the algorithm excludes the hairpin and ends the helix at the preceding stem. Hairpins have no assigned bend angle. The system reports a single-segment canonical stem-loop only if its hairpin passes the stacking test.

In the second stage, geometric validation limits the bending of the assembled helix. For each accepted connector, the algorithm calculates the local bend angle between the helical axis vectors of the two flanking stems, obtained from X3DNA. The algorithm incorporates a connector into the growing helix only if this local bend angle is below or equal to the 50° threshold (see Section 3.3.1). In addition, the algorithm continuously tracks the global bend angle between the helical axis of the initial stem and that of the most recently added stem. If this global angle exceeds the threshold, the algorithm terminates the extension and finalizes the current assembly. Rather than being discarded, the non-qualifying connector becomes the start of a new helix candidate. This mechanism enables the detection of consecutive quasi-linear segments within a single RNA chain and ensures that each resulting n-segment extended helix forms a structurally coherent unit.

### 3.5 Junction analysis

We classify junctions topologically by the presence of *n* ≥ 3 annotated loop strands. A compactness check enforces structural integrity by inspecting PDB author numbering and insertion codes and treating nonconsecutive numbering as a gap. This check excludes junctions that contain sequence gaps or non-RNA residues. To characterize tertiary stabilization within the junction, we analyze coaxial stacking using a three-step graph method.

First, we construct a stacking graph where nodes represent nucleotides and edges represent base-stacking interactions. Graph traversal is restricted to loop nucleotides and terminal positions of flanking stems, which prevents spurious paths through unrelated regions of the structure.

Second, a breadth-first search algorithm identifies all continuous stacking paths between the terminal base pairs of any two stems that radiate from the junction core.

Third, we filter candidate coaxial pairs to retain only pairs with an inter-stem bend angle below or equal to 50°, computed from X3DNA helical axis vectors (see Section 3.3.1). We rank the remaining pairs in ascending order of bend angle and apply a greedy selection algorithm to ensure that no stem participates in more than one coaxial stack. We exclude junctions without a geometrically valid coaxial pair from further analysis.

### 3.6 Calculation of structural parameters in extended elements

For complex branched topologies, the calculation of structural parameters is strictly restricted to the continuous path formed by coaxially stacked stems. The total nucleotide count includes exclusively the residues of these stacked flanking stems and the specific loop path connecting them. Consequently, non-canonical base pairs are counted only if both interacting nucleotides belong to this defined continuous path. Any pairs or residues associated with unstacked, radiating stems are explicitly excluded from the extended helix statistics. Finally, all non-canonical interactions are deduplicated to prevent higher-order contacts, such as base triples, from artificially inflating the final pair count.

### 3.7 Visualization

We represent structures through combined 2D and 3D visualizations. A dual-palette color scheme maintains consistency across both views by distinguishing helix components from junction cores. For 3D rendering, automated PyMOL scripts isolate each structural element using queries based on chain identifiers, residue numbers, and insertion codes. Each script sets viewing parameters, applies cartoon and stick representations, and centers the camera on the target element.

For 2D topology, the pipeline exports JSON files compatible with the VARNA visualization engine. It maps internal serial indices back to standard PDB author numbering to preserve biological context. Canonical Watson-Crick base pairs are derived directly from the dot-bracket secondary structure, whereas non-canonical interactions follow the Leontis-Westhof classification [43]. Lines connecting non-consecutive stacked nucleotides across loop regions show coaxial stacking paths and tertiary continuity. A thread pool executes the pipeline in parallel by batching VARNA tasks across all structural elements, and the pipeline converts the resulting JSON data into SVG images.

## 4 Conclusions

RNAbridge provides, to our knowledge, the first database to extract extended non-canonical helices and coaxially stacked multi-way junctions as objects with their own identifiers, statistics, and renderings. Applying a geometry- and stacking-based pipeline to RNA-containing PDB entries, the database catalogs 153,148 structural elements, of which 7,453 form the representative view. In this view, 62.0% of the 4,998 angle-analyzed elements have bends below 20°, and three-way junctions are the most common branched topology (861 of 1,862 junction arrangements).

Beyond its scale, RNAbridge’s value is illustrated by targeted case studies, from resolving the non-canonical base-pairing network that unifies the five segments of ribosomal helix 44 into a single quasi-continuous helix to directly visualizing the stacking interactions that define Lescoute and Westhof’s Family C three-way junction topology. By combining synchronized 2D and 3D representations, rich searchable metadata, and weekly automatic updates in an accessible web interface, RNAbridge offers RNA structural biologists and 3D modelers a practical tool for exploring the tertiary architecture that canonical secondary-structure representations inherently miss. Moreover, it establishes a foundation for future work extending this framework to other classes of higher-order RNA structures. In the future, these data may support machine learning models [44] for predicting coaxiality from multiloop sequences.

The catalog remains subject to PDB deposition bias: reported frequencies describe available structures rather than biological abundance. The representative view reduces repeated sampling but does not entirely remove this bias. Because pseudoknot stems are not cataloged as independent stems and the terminal-hairpin test is restricted to the loop, pseudoknotted stem-loops may not qualify as single-segment entries. Extending the catalog to pseudoknot-stabilized hairpins is left for future work.

## CRediT authorship contribution statement

**Damian Zakrzewski**: Software, Visualization, Writing - original draft. **Maciej Antczak**: Conceptualization, Methodology, Validation, Writing - original draft, Writing - review and editing. **Tomasz Zok**: Conceptualization, Funding acquisition, Methodology, Project administration, Software, Validation, Writing - original draft, Writing - review and editing.

## Data availability

The web server is publicly accessible, and its source code is available. All data used for the experiments were sourced from public resources. Web server: https://rnabridge.cs.put.poznan.pl. Source code: https://github.com/onidd/RNAbridge [45]. License: MIT. All counts reported in this paper refer to the November 8, 2026 release of the database. The current source-code release is archived at https://doi.org/10.5281/zenodo.20428883.

## Acknowledgements

This work is supported by funds from the National Science Center, Poland [2023/51/D/ST6/01207 to TZ] and statutory funds of the Institute of Computing Science, Poznan University of Technology.

