## Supplementary Materials for "RNAbridge: a database of extended non-canonical helices"

Damian Zakrzewski[1], Maciej Antczak[1,2], and Tomasz Zok *[1]

[1]Institute of Computing Science, Poznan University of Technology, ul. Jacka Rychlewskiego 1, 61-131, Poznan, Poland

[2]Department of Structural Bioinformatics, Institute of Bioorganic Chemistry PAS, Noskowskiego 12/14, 61-704, Poznan, Poland

All counts refer to the November 8, 2026 release. Tables S1 and S2 give nucleotide-count distributions for the full catalog and the representative view (labeled "non-redundant" in the web interface), respectively. Representative-source selection reduces repeated sampling but does not establish non-redundancy of extracted local elements. Table S3 gives the full-catalog bend-angle distribution at the 50° cutoff, and Table S4 reports the sensitivity of retained counts to this cutoff. In Tables S1 and S2, 1-seg entries are canonical stem-loops whose terminal hairpins pass the stacking-continuity test, not a census of all canonical stem-loops. Multi-segment helices are grouped by stem count and junctions by order; "+" denotes the indicated value or higher.

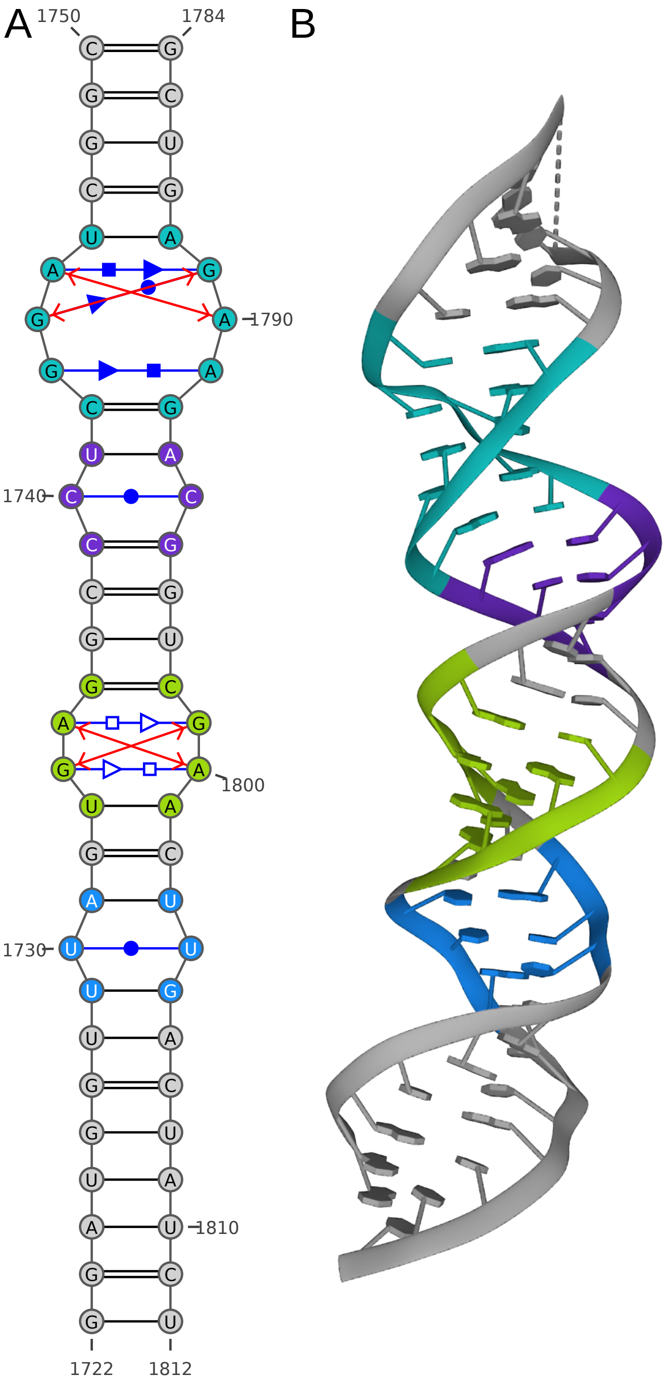


Figure S1: Helix 44 fragment for the structure of the human 48S initiation complex in closed state (PDB id: 7QP7). (A) 2D diagram, (B) 3D visualization.

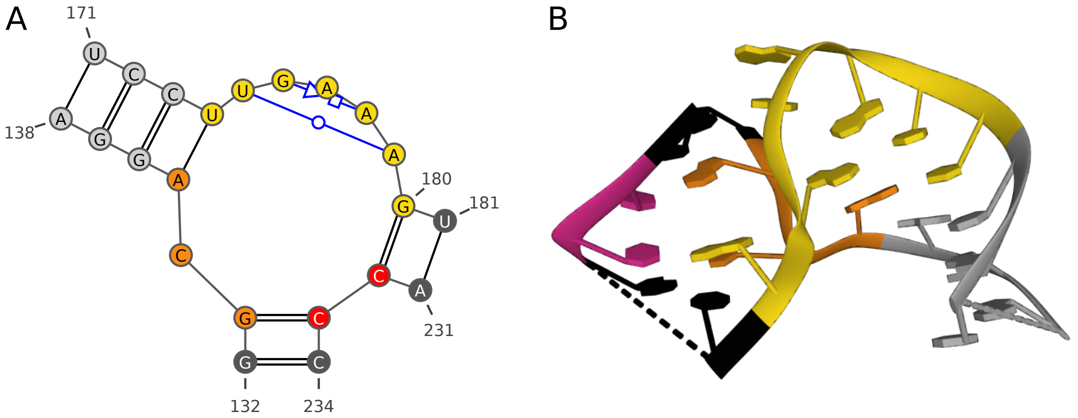


Figure S2: Three-way junction of Family C type from the crystal structure of the specificity domain of Ribonuclease P RNA (PDB id: 1NBS). (A) 2D diagram, (B) 3D visualization.

Table S1: Nucleotide-count distribution of the full catalog (153,148 elements).

| Unique nucleotides / Type | 1-seg | 2-seg | 3-seg | 4-seg+ | 3-way | 4-way | 5-way | 6-way+ | Total |
|---|---|---|---|---|---|---|---|---|---|
| [0, 10) | 20,677 | 1,089 | – | – | 430 | 15 | 3 | – | 22,214 |
| [10, 20) | 24,420 | 31,942 | 403 | – | 17,446 | 8,584 | 2,446 | 1,231 | 86,472 |
| [20, 30) | 4,078 | 20,745 | 2,826 | 21 | 1,411 | 6,019 | 148 | 846 | 36,094 |
| [30, 40) | 44 | 1,910 | 1,780 | 239 | 1,221 | 143 | 263 | 1,656 | 7,256 |
| [40, 50) | 2 | 537 | 171 | 178 | 35 | 7 | 13 | 2 | 945 |
| [50, 60) | 1 | 11 | 26 | 14 | – | 8 | – | – | 60 |
| [60, 70) | 1 | – | 3 | 89 | – | 1 | – | – | 94 |
| [70, 80) | 1 | – | – | 7 | – | – | – | – | 8 |
| [80, 90) | 1 | – | – | 1 | – | – | – | – | 2 |
| [90, 100) | 1 | 2 | – | – | – | – | – | – | 3 |
| Total | 49,226 | 56,236 | 5,209 | 549 | 20,543 | 14,777 | 2,873 | 3,735 | 153,148 |

Table S2: Nucleotide-count distribution of the representative view (7,453 elements).

| Unique nucleotides / Type | 1-seg | 2-seg | 3-seg | 4-seg+ | 3-way | 4-way | 5-way | 6-way+ | Total |
|---|---|---|---|---|---|---|---|---|---|
| [0, 10) | 908 | 76 | – | – | 17 | 3 | – | – | 1,004 |
| [10, 20) | 1,290 | 1,604 | 25 | – | 649 | 364 | 86 | 36 | 4,054 |
| [20, 30) | 244 | 1,021 | 148 | 3 | 138 | 366 | 26 | 40 | 1,986 |
| [30, 40) | 11 | 87 | 106 | 14 | 55 | 27 | 1 | 40 | 341 |
| [40, 50) | 1 | 14 | 10 | 11 | 2 | 4 | 4 | – | 46 |
| [50, 60) | – | 6 | 1 | – | – | 4 | – | – | 11 |
| [60, 70) | 1 | – | 2 | 6 | – | – | – | – | 9 |
| [70, 80) | – | – | – | 1 | – | – | – | – | 1 |
| [80, 90) | – | – | – | 1 | – | – | – | – | 1 |
| [90, 100) | – | – | – | – | – | – | – | – | – |
| Total | 2,455 | 2,808 | 292 | 36 | 861 | 768 | 117 | 116 | 7,453 |

Table S3: Bend-angle distribution of the full catalog at the 50° cutoff (103,922 elements). The 49,226 single-segment entries are excluded because no inter-helical angle can be computed.

| Angle / Type | Multi-segment Helices | | | Junction-based Arrangements | | | | Total |
|---|---|---|---|---|---|---|---|---|
| | 2-seg | 3-seg | 4-seg+ | 3-way | 4-way | 5-way | 6-way+ | |
| [0°, 5°) | 3,697 | 59 | 37 | 632 | 1,337 | 639 | 26 | 6,427 |
| [5°, 10°) | 8,755 | 227 | 86 | 2,780 | 2,376 | 1,006 | 80 | 15,310 |
| [10°, 15°) | 9,822 | 495 | 78 | 4,955 | 2,604 | 267 | 298 | 18,519 |
| [15°, 20°) | 10,263 | 865 | 36 | 4,057 | 1,731 | 425 | 797 | 18,174 |
| [20°, 25°) | 9,051 | 819 | 41 | 2,519 | 1,525 | 233 | 379 | 14,567 |
| [25°, 30°) | 6,517 | 773 | 33 | 1,728 | 1,594 | 144 | 1,617 | 12,406 |
| [30°, 35°) | 4,507 | 979 | 78 | 1,178 | 1,228 | 85 | 399 | 8,454 |
| [35°, 40°) | 2,336 | 495 | 93 | 1,653 | 1,575 | 31 | 93 | 6,276 |
| [40°, 45°) | 964 | 345 | 59 | 610 | 538 | 24 | 31 | 2,571 |
| [45°, 50°] | 324 | 152 | 8 | 431 | 269 | 19 | 15 | 1,218 |
| Total | 56,236 | 5,209 | 549 | 20,543 | 14,777 | 2,873 | 3,735 | 103,922 |

Table S4: Sensitivity of the full-catalog element counts to the bend-angle cutoff. Counts are re-derived at each threshold; 1-seg entries are unaffected and excluded. The threshold affects helix assembly, so the 0–50° bins are not identical across runs. Change is relative to the 50° count; the 90° versus 70° change is +0.23%.

| Threshold | 2-seg | 3-seg | 4-seg+ | 3-way | 4-way | 5-way | 6-way+ | Total | Change vs 50° |
|---|---|---|---|---|---|---|---|---|---|
| 50° | 56,236 | 5,209 | 549 | 20,543 | 14,777 | 2,873 | 3,735 | 103,922 | – |
| 70° | 55,708 | 5,576 | 564 | 21,816 | 15,549 | 2,949 | 3,760 | 105,922 | +1.9% |
| 90° | 55,735 | 5,578 | 567 | 21,871 | 15,655 | 2,996 | 3,765 | 106,167 | +2.2% |
